# Binning unassembled short reads based on k-mer covariance using sparse coding

**DOI:** 10.1101/599332

**Authors:** Olexiy Kyrgyzov, Vincent Prost, Stéphane Gazut, Bruno Farcy, Thomas Brüls

**Affiliations:** CEA Genoscope, Evry, France; CEA LIST, Gif-sur-Yvette, France; Bull Technologies, Les Clayes-sous-Bois, France

**Keywords:** Metagenomics, Sequence binning, Sparse coding

## Abstract

Sequence binning techniques enable the recovery of a growing number of genomes from complex microbial metagenomes and typically require prior metagenome assembly, incurring the computational cost and drawbacks of the latter, e.g. biases against low-abundance genomes and inability to conveniently assemble multi-terabyte datasets.

We present here a scalable pre-assembly binning scheme (i.e. operating on unassembled short reads) enabling latent genomes recovery by leveraging sparse dictionary learning and elastic-net regularization, and its use to recover hundreds of metagenome-assembled genomes, including very low-abundance genomes, from a joint analysis of microbiomes from the LifeLines-Deep population cohort (n=1135, > 10^10^ reads).

We showed that sparse coding techniques can be leveraged to carry out read-level binning at large scale, and that despite lower genome reconstruction yields compared to assembly-based approaches, bin-first strategies can complement the more widely used assembly-first protocols by targeting distinct genome segregation profiles. Read enrichment levels across six orders of magnitude in relative abundance were observed, indicating that the method is able to recover genomes consistently segregating at low levels.

## 1 Introduction

Metagenomic shotgun sequencing has dramatically increased our appreciation of the intricacies of microbial systems, whether sustaining biogeochemical processes or underlying health status of their hosts. Several limitations, including sequencing errors, strain-level polymorphism, repeat elements and inequal coverage, among others, concur however to yield fragmented metagenome assemblies, requiring post-processing in order to cluster (bin) assembled fragments into meaningful biological entities, ideally strain-resolved genomes.

The advent of reasonably efficient sequence binning techniques, often exploiting a coverage covariance signal across multiple samples, allowed the field of metagenomics to move toward more genome-centric analyses [7], and recently thousands of so-called metagenome-assembled genomes (MAGs) have been reported, both from environmental sources and human surfaces or cavities [2, 28, 30, 31]. The vast majority of these MAGs have been produced by post-assembly binning approaches, i.e. operating on sequence contigs assembled on a sample by sample basis. Though highly successful, such methods are nevertheless “inherently biased towards the most abundant organisms, meaning consistently less abundant organisms may still be missed” (quoted from ref [2]). For example, although thousands of MAGs were reconstructed from more than 1500 public metagenomes in the remarkable study [30], over 93% of these MAGs had an average coverage of more than 10x in at least one of the samples analyzed. The high proportions of phylogenetically unassigned reads typical in medium to high complexity metagenomes is another consequence of this limitation [13].

Even though the ecological or community-level importance of rare species is a matter of debate, there are both theoretical and empirical observations supporting the notion that rare organisms can substantially contribute to community-level behavior and resilience, hence represent valuable targets for genome recovery. Theoretical modeling of microbial trade of diffusible goods [18] have for example highlighted an apparent paradox (called “curse of increased efficiency” by the authors of ref [18]), where one bacterial species becomes rarer in the population despite becoming fitter and more efficient at producing a key metabolic resource. This situation is provoked by metabolic interdependencies that can evolve via trade in microbial consortia, and that can lead to low-abundance organisms becoming essential for a faster growth rate of the community. On the other hand, several empirical studies have documented the ecosystem-level relevance of rare bacteria (see ref [16] for a review), for example ref [17] makes a case for the role of “ultrarare” bacteria in ecosystem-level productivity, and ref [5] highlight the role of some low abundance bacteria in driving termite’s hindgut bacterial community composition.

Considering that global metagenome assembly (or cross-assembly) is currently unpractical to recover low abundance genomes or complex microbial consortia from terabytes of data, we decided to investigate a “bin first and assemble second” paradigm that could make the assembly problem more tractable by targeting lower complexity sequence subsets (bins). Binning unassembled reads is however more computationally demanding, as the number of raw sequences is typically orders of magnitude higher than the number of assembled contig sequences.

Even though the dominating paradigm nowadays is assembly-first binning, it is worth noting that the first sequence binning methods reported, like AbundanceBin [41] and MetaCluster [42], operated at the read level. This shift towards contig binning was mainly driven by the increase in data throughput, as the first read-level binning methods were designed at the time of 454 (Roche) and even Sanger sequencing (both providing longer reads) to process individual samples. They were thus not designed to scale to large multi-sample terabase-sized short read datasets. In this perspective, assembly can be viewed as a pre-processor to reduce the computational burden of binning.

A pioneering pre-assembly binning scheme [11] was proposed a couple of years ago, with the read partitioning problem formulated by analogy to the latent semantic analysis (LSA) technique widely used in natural langage processing (NLP). The core idea to view metagenomes as linear mixtures of genomic variables can lead to read clustering formulations based on the deconvolution of latent variables (“eigengenomes”) driving the k-mer (subsequences of length k) abundance covariance across samples. The raw sequence data is first summarized in a sample by k-mer occurrence matrix (analogous to term-document matrices in NLP), approximating the abundance of k-mers across samples. Matrix decomposition techniques can then be used to define two sets of orthogonal latent vectors analogous to principal components of sample and sequence space. The large memory requirements incurred by the factorisation of large abundance matrices naturally drove ref [11] toward a rank-reduced singular value decomposition (SVD), for which efficient streaming libraries [33] enable a parallel processing of blocks of the abundance matrix by updating the decomposition iteratively. Clusters of k-mers can then be recovered by an iterative sampling and merging heuristic that samples blocks of eigen k-mers from the right singular vectors matrix until an arbitrary portion (about 0.4% in ref [11]) of the latter has been covered. This heuristic is however acknowledged as a significant hindrance, the authors calling for “more sophisticated methods [are needed] to computationally discover a natural clustering” (quoted from ref [11]).

We describe here a pre-assembly binning method based on sparse dictionary learning and elastic-net regularization that exploits sparsity and non-negativity constraints inherent to k-mer count data. This sparse coding formulation of the binning problem can leverage efficient online matrix factorization techniques [25] and scales to very large (terabyte-sized) k-mer abundance matrices; it also bypasses the aforementioned problematic k-mer clustering heuristic, removes interpretability issues associated with the SVD (e.g. the physical meaning of negative contributions), and is able to enrich sequences from a given genome across six orders of magnitude in relative abundance (see section “Recovery of very low-abundance genomes” thereafter).

## 2 Results

We describe in the following section some results obtained with the proposed binning scheme based on the modeling of data vectors as sparse linear combinations of basis elements (sparse coding [25]).

We will start with a preliminary experiment illustrating the ability of read binning to recover a target genome whose sequences segregate at levels too low to yield any kilobase-sized fragment by assembly in any single sample, hence would not be recoverable by assembly-first approaches. We will then describe results from a direct comparison of assembly-first versus bin-first methods that illustrate the complementarity of the two approaches in terms of the profiles of genomes recovered. The next subsection describes a comparison of the sparse coding based bin-first approach with a state of the art read-binning method. The next subsections will describe strain separation results obtained with the new method, document its scalable behavior, and its ability to enrich rare sequences, thereby enabling the recovery of low abundance genomes. We will conclude with a discussion of some important limitations of the method and consider some of its potential applications.

### 2.1 Read-level binning can recover low abundance genomes that escape assembly-first protocols

We devised an experiment to illustrate a situation where assembly-first approaches are not able to recover a target genome -because target genome sequences are too low in number in any single sample-whereas a bin-first approach is successful at it. The experimental setup involved distributing a very low number of short reads (100 paired reads) randomly sampled from a target genome (a 10 kbp plasmid) into 14 samples containing each a background of 20000 unrelated bacterial sequences (4 further samples contained only background sequences with no read from the target genome at all). As no single kilobase-sized fragment could be recovered by assembling the sequences from each sample individually, this precluded the application of assembly-first methods (e.g. contig binning methods like metabat [19, 20] require >= 1500 bp sequences as input). On the other hand, circa 90% of the reads originating from the target genome could be segregated in a single cluster/bin using our read binning pipeline (Supplementary Table 1), leading to the complete recovery of the target genome in a single contig after assembly (Methods).

### 2.2 Bin-first and assembly-first strategies recover distinct and complementary genome sets

A second experiment aimed at directly comparing the genome recovery yield of assembly-first versus bin-first strategies on a real-life dataset. We selected the raw sequence data from 18 (randomly chosen) individuals of the LifeLinesDeep cohort [43], and either assembled these individually (i.e. on a sample by sample basis) with metaSPAdes (v3.13.0) followed by contig binning across samples with the MetaBat2 adaptive algorithm [19], or clustered the raw reads using our read-level binning pipeline, followed by metaSPAdes assembly of the resulting partitions/bins.

Fourteen nearly (>90%) complete and uncontaminated (<5%) genomes were recovered using the assembly-first approach, versus seven using the bin-first method. Interestingly, the two genome sets were disjoint, with no complete genome recovered by both approaches. Among the 14 genomes recovered by the assembly-first approach, three were not represented in the set of 164 MAGs recovered from the analysis of the entire cohort using the bin-first protocol. More surprisingly, only three out of the seven complete genomes retrieved by our bin-first pipeline from the analysis of 18 samples were represented among the complete or nearly complete MAGs identified from the full cohort analysis, indicative of a lack of stability of the algorithm that we relate to bin fragmentation provoked by extensive strain-level variation across the samples (see Discussion).

The surprising lack of overlap between the two genome sets in this experiment is not attributable to fundamental differences in abundance levels between the genomes recovered by the two approaches, as in both cases the genome bins could be directly aligned to individual sample assemblies, i.e. the genomes recovered using both approaches were of sufficiently high coverage to yield relatively large contigs in the assemblies of individual samples. We assessed potential differences between the distributions of binned genome sequences across the samples, which highlighted distinct patterns for the two approaches, with the genomes identified by the bin-first approach aggregating sequences from a larger number of samples (and harboring a higher number of contigs per genome bin on average) (Figure 1).

**Fig. 1:**
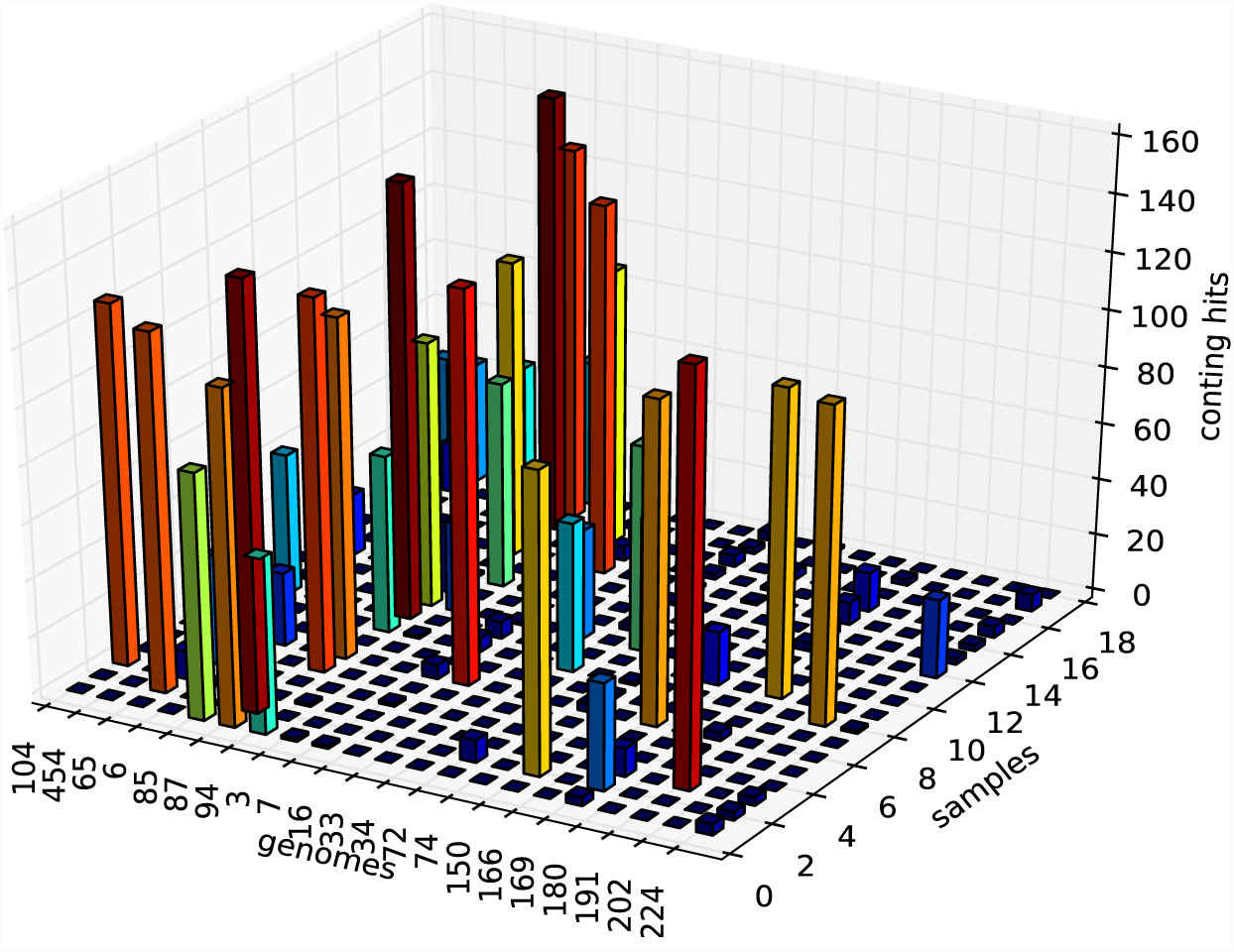
Sample origins of the sequences aggregated into genome bins (displayed by their genome identifier on the x-axis) using our bin-first method (first seven genomes (104 to 94) on the left) versus assembly-first binning using metabat2 (fourteen rightmost (3 to 224) genomes). Genomes retrieved by the bin-first method aggregate sequences from a larger number of samples.

Thus, in the present experiment, the assembly-first approach targeted genomes reaching high abundance in a limited number of samples, for which the weaker abundance covariation signal probably hampered the bin-first approach. Consistent with this view, sequences from genomes produced through the assembly-first approach were frequently located in large (dozens of Mbp in size) and unresolved partitions computed by read-level binning (see Discussion).

On the other hand, we should keep in mind that the number of samples (18) used in this experiment is relatively low. Related approaches based on abundance-covariance, like Concoct [3] or LSA [11] among others, require a higher number of samples to achieve best performance (about 50 samples for the former and between 30 to 50 for the latter).

Despite these limitations, the fact that the bin-first approach was able to recover a significant number of complete genomes not identified by the assembly-first approach illustrates the complementarity of the two strategies.

### 2.3 Enhanced accuracy of sparse coding based read binning versus state of the art read binning

Besides the two pioneering read binning methods already mentioned (AbundanceBin [41] and MetaCluster [42]), we could also mention CompostBin [9], which is a PCA-based read level binning algorithm that was designed and tested on Sanger reads. BiMeta [38] and MetaProb [14] are other tools that operate at the read-level, but describe themselves as “assembly-assisted”, meaning they rely on the detection of read overlaps. BiMeta was tested on 454 reads simulating bacterial communities of a dozen of different genomes at most and on the Acid Mine Drainage dataset of ref [37], which is of low complexity and consists in Sanger reads. MetaProb shares some principles with BiMeta: it is also “assembly-assisted” and was tested on the same low-complexity synthetic datasets as the latter. The authors also tested their method on a real microbiome sample consisting in 43 million reads, but only after filtering the latter down to 2 million reads. Thus, all the above methods were designed to operate on a individual samples, at a time were scalability issues were less acute. Moreover, with the exception of AbundanceBin which exploits a coverage signal extracted from unique k-mers, the other methods are better described as composition-based, using a nucleotide composition signal measured from short k-mers (typically of length 4 or 5).

We developed our method with scalability in mind, as we wanted it to be able to process on the order of 10^10^ short reads and to be able to process increasingly larger multi-sample datasets by simply stacking additional computing resource. In this respect, there is only one competing method left, Latent Strain Analysis [11], that is both scalable and designed to operate on unassembled short reads from a large number of samples.

To evaluate our method, we first compared its read clustering accuracy (measured in terms of precision, recall and F-value metrics, see Methods) with that of the original LSA method by using previously described benchmark datasets [15] (downloadable from http://www.genoscope.cns.fr/SCdata/vc50/), for which read to genome assignments were known (ref [15] and Methods). The results from these experiments are summarized in Table 1, and show improved accuracy of the sparse-coding framework over both the original LSA and a naive k-means algorithms.

**Table 1:** Binning accuracy estimates: LSA refers to the original algorithm of ref [11], with a cosine similarity threshold of 0.7 as recommanded by the authors, k-means refers to a direct clustering of the columns of the abundance matrix, with the number of clusters set to 1000 (equal to the number of components for the sparse decomposition), see main text and Methods.

|  | K-means | LSA | Sparse Coding |
| --- | --- | --- | --- |
| Precision | 0,52 | 0,58 | 0,72 |
| Recall | 0,63 | 0,64 | 0,82 |
| F-Value | 0,57 | 0,61 | 0,77 |

### 2.4 Partial strain separation

The counting and indexing of k-mers in fixed memory is achieved by locality sensitive hashing (Methods). By design, locality sensitive hash functions increase the probability of collision for related items [8]. On one hand, this provides a convenient way to handle sequencing errors. On the other hand, the occurrence in natural environments of multiple strains from the same species (the so-called species pangenome) could lead to artefactual k-mer merging and potential overlap between distinct genomic partitions. This represents an issue potentially exacerbated by the inter-sample read aggregation process.

To assess the behavior of the method in the presence of extensive pangenomic (i.e. strain-level) variation, we quantified its ability to separate closely related (up to 99.96% average nucleotide identity (ANI), Table 2) strains that were deliberately included in the genome mixtures from the virtual cohort used in the test experiments.

**Table 2:**
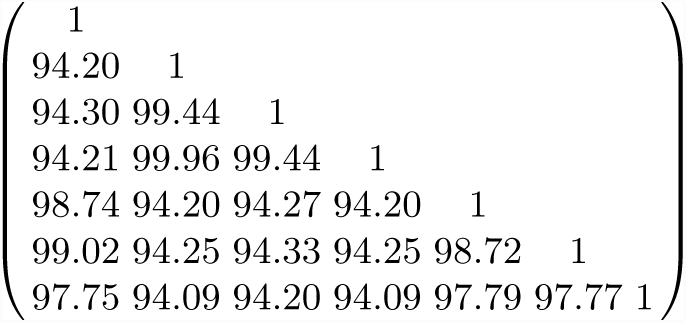
Average Nucleotide Identity (ANI) between the *B. amyloliquefaciens* strains used in the strain separation experiment illustrated in the left panel of Figure 2 (see main text and Methods)

The two panels of the Figure 2 illustrate two practical examples of partial strain separation achieved with the method. The left panel illustrates a partial separation of 7 strains from the *Bacillus amyloliquefaciens* species (whose ANI ranged from 94.18 to 99.96, Table 2), while the right panel shows similar results for 8 strains of the *Sulfolobus islandicus* species (whose ANI ranged from 97.84 to 99.59). As the genomic origin of each read is known in the virtual cohort dataset, these plots show, for each strain (represented by a horizontal line), the distribution of its reads among the full set of clusters/bins generated by the pipeline (and arbitrarily ordered along the x-axis). The left panel illustrates that the 7 strains from the *B. amyloliquefaciens* species are mostly separated into two groups according to whether their main cluster is located near x-coordinate 220 or x-coordinate 500. The right panel on the other hand shows that the 8 strains of the *S. islandicus* species share a common “core” cluster (located near the origin), while variable portion of their genomes are segregated into distinct “variable” clusters.

**Fig. 2:**
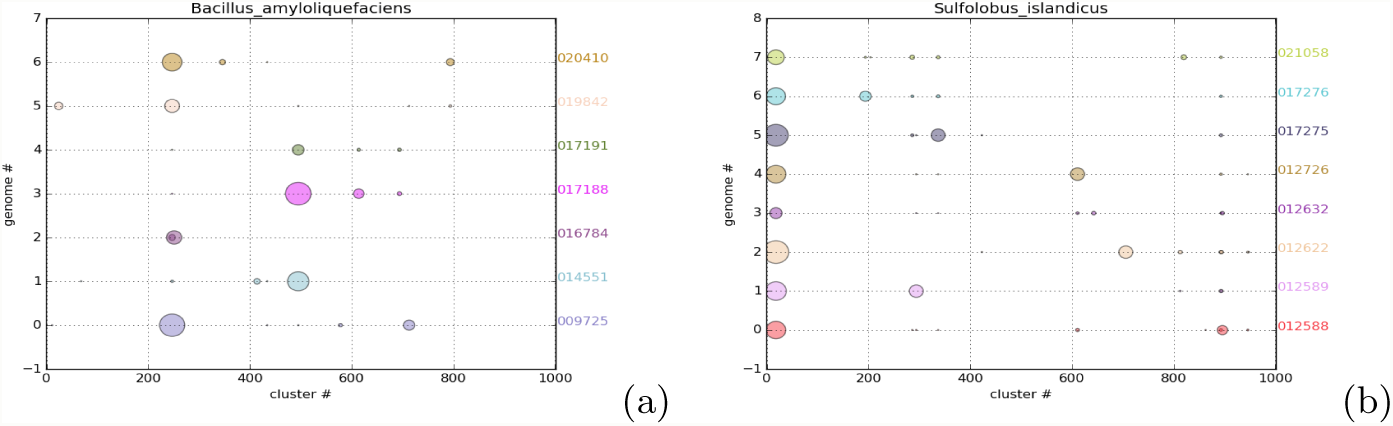
Partial resolution of species pangenomes. x-axis: partition identifier, y-axis: horizontal axes correspond to different strains from the same species (left: *B. amyloliquefaciens* strains, right: *S. islandicus* strains). Circle area is proportional to the number of reads from a given strain assigned to the given partition. The left panel illustrates the partial separation of seven strains in two distinct partitions. The right panel illustrates the differential segregation of the core (at the left of the figure) and variable portions of the species pangenome.

Overall, this analysis makes apparent a partial separation of closely related strains (left panel), as well as the differential segregation of the core (i.e. the genome fraction that is shared between all the strains of a species) and variable portions of the species pangenomes (right panel). In practice, some level of strain mix-up is probably inherent to the inter-sample read aggregation process, and approaches based on sample by sample assembly limit the risk of strain mixing, but at the expense of focusing on those genomes that reach high-coverage (around 10x). Our approach aimed at relaxing the latter constraint, but by doing so through the aggregation of lower abundance reads across samples, it becomes vulnerable to extensive strain-level variation. Dealing with this problem is the focus of future research, e.g. a possible workaround could be to carry out a “soft-clustering” by allowing “core” sequences to belong to more than one “variable” cluster.

### 2.5 Sensitivity and scalability on real-life data

By scalability, we refer to the ability of the method to adapt to order of magnitude change in the input (raw reads), and its ability to maintain its functionality and performance under high demand (i.e increasingly higher data volumes).

To assess the sensitivity and scalability of the sparse coding method, we applied it to a real world dataset of over 10^10^ reads (about 10 terabytes of raw sequence data) derived from 1135 gut microbiomes of healthy Dutch individuals from the LifeLinesDeep cohort [43]. The pre-assembly binning of the cohort’s reads resulted in 983 partitions, which were then assembled individually using the Spades engine [4] (Methods). The distribution of assembly sizes is shown in Figure 3, making apparent that the vast majority of partitions are bacterial-genome sized (i.e. in the 2-5 Mbp range). A few dozens of coarse-grained partitions harboring unresolved genomes makes up the right tail of the distribution. As a direct read to genome mapping is not available for real life metagenomes, we assessed clustering performance by quantifying the genomic homogeneity and completeness of the resulting partitions based on the occurrence pattern of universal single-copy markers using the checkm toolkit [29]. A summary of completion and contamination statistics of the genome-resolved partitions is presented in Table 3, while another facet of the homogeneity of reconstructed genomes is displayed in the left panels of Figure 5.

**Table 3:** Genome completion and contamination statistics of assembled partitions/bins, see main text and Methods.

| Classification | Completeness | Genomes (Bins) | Contamination |
| --- | --- | --- | --- |
| Nearly Complete | >90% | 14 | $\leq 5\%$ |
| Substantial | >70% to $\leq 90\%$ | 53 | $\leq 5\%$ |
| Moderate | > 50% to $\leq 70\%$ | 97 | $\leq 5\%$ |
| Partial | $\leq 50\%$ | 724 | $\leq 5\%$ |
| Unresolved | >100% | 95 | > 5% |

**Table 4:** Cluster assignments of reads from a target genome versus background (unrelated) reads. Nearly all the 2800 reads from the target genome segregating at low levels in the samples (100 paired-reads per sample in 14 samples, none in the remaining samples) are binned in a single partition using our bin-first pipeline, leading to the complete genome after assembly. No kilobase-sized contig could be assembled from any individual sample, making the assembly-first protocol inoperable (see main text).

| cluster id | # of spiked reads | # of reads | ratio |
| --- | --- | --- | --- |
| 0 | 2782 | 5106 | 0.544849 |
| 1 | 0 | 6164 | 0.000000 |
| 2 | 0 | 7284 | 0.000000 |
| 3 | 0 | 4352 | 0.000000 |
| 4 | 0 | 5774 | 0.000000 |
| 5 | 6 | 6594 | 0.000910 |
| 6 | 0 | 4498 | 0.000000 |
| 7 | 0 | 5134 | 0.000000 |
| 8 | 0 | 6380 | 0.000000 |
| 9 | 0 | 7206 | 0.000000 |
| 10 | 0 | 5490 | 0.000000 |
| 11 | 0 | 6220 | 0.000000 |
| 12 | 0 | 6574 | 0.000000 |
| 13 | 0 | 6252 | 0.000000 |
| 14 | 0 | 6230 | 0.000000 |
| 15 | 0 | 6624 | 0.000000 |
| 16 | 0 | 5600 | 0.000000 |
| 17 | 8 | 6704 | 0.001193 |
| 18 | 2 | 6802 | 0.000294 |
| 19 | 2 | 67812 | 0.000029 |

**Fig. 3:**
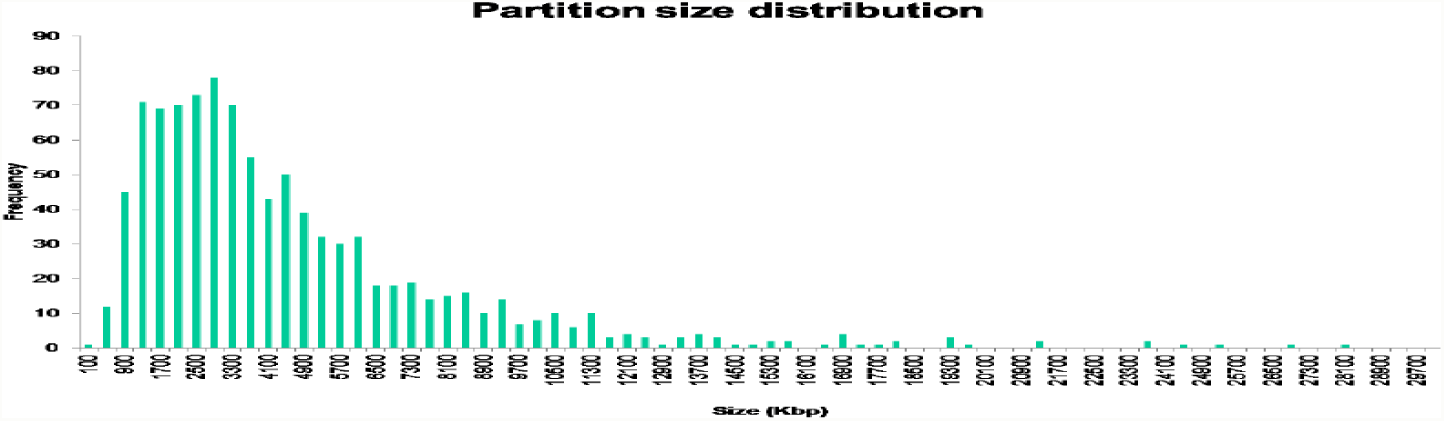
Distribution of assembled bin sizes. x-axis: assembled partition size (in kbp), y-axis: partition frequency

**Fig. 4:**
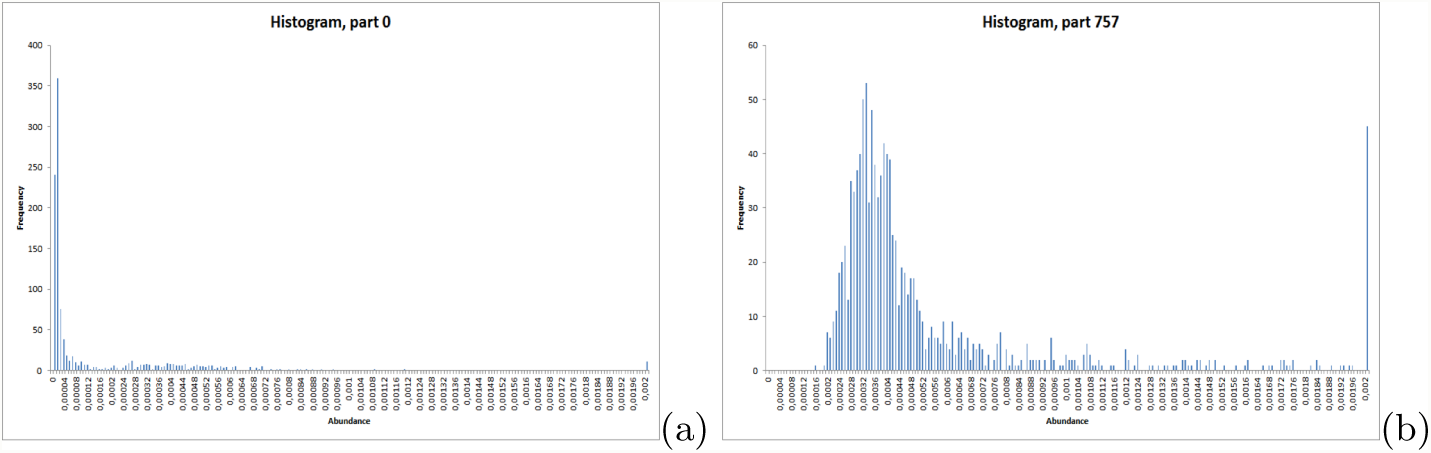
Enrichment histograms displaying the fraction of raw reads contributed by each sample to two distinct genome-resolved bins. x-axis: read abundance of partition 0 (left) and partition 757 (right); y-axis: sample frequency (among the 1135 samples). Different situations are illustrated: a relatively high proportion of reads can be contributed by a small subset of individuals (a few dozens, corresponding to the rightmostest peak for the genome-resolved bin shown in panel b), while the left panel illustrates that substantial (i.e. ≥70% complete) genomes of low-abundance organisms can also be recovered by aggregating only a few thousands reads per sample across the full cohort.

**Fig. 5:**
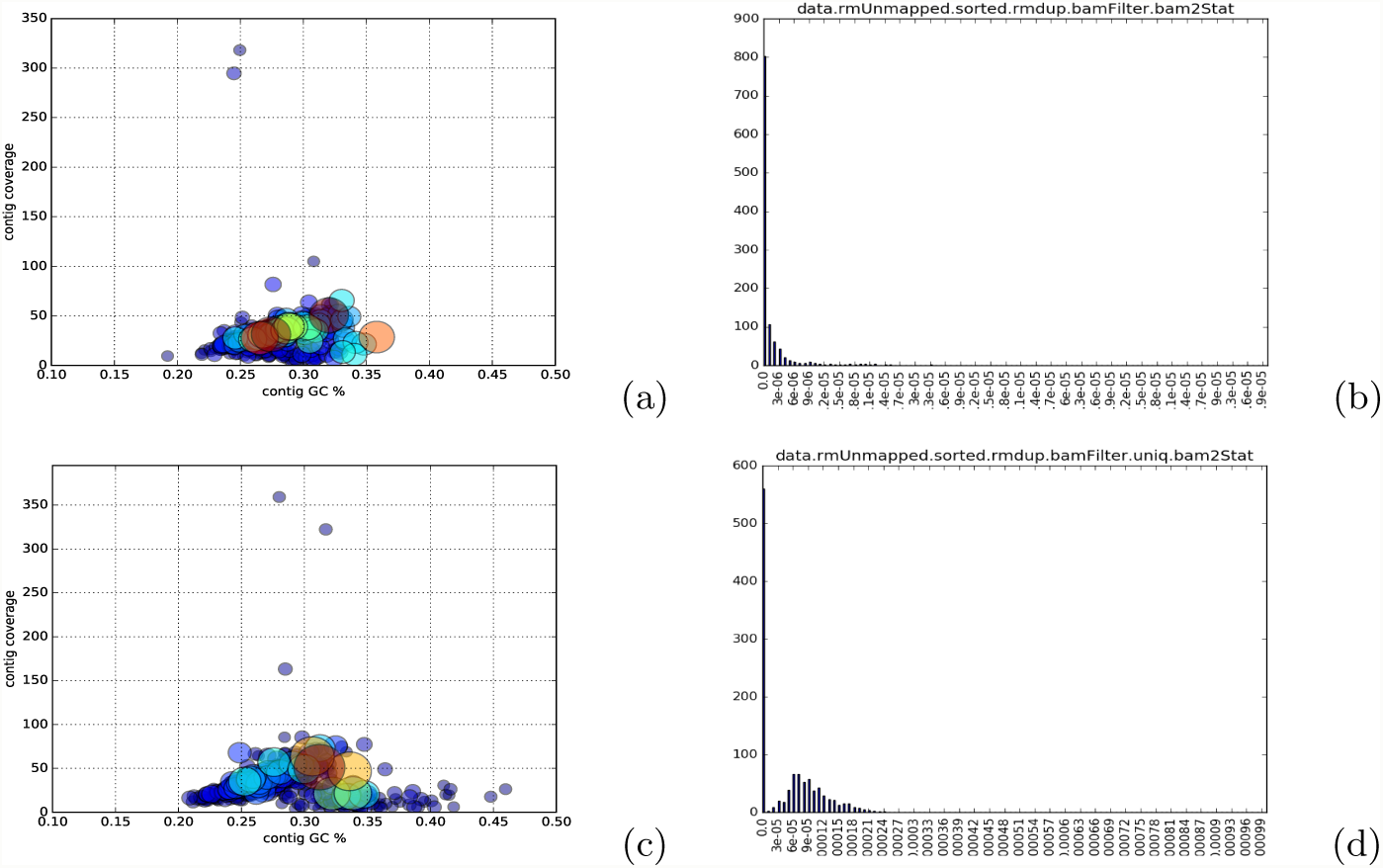
Left panels (a,c): GC-coverage plots (x-axis: contig GC%; y-axis: contig coverage) illustrating the homogeneity of two assembled bins (top: bin 470 (70% complete, 4.8% contamination) bottom: bin 766 (70% complete, 3.5% contamination)) corresponding to two unclassified Firmicutes genomes of low-abundance, whose enrichment histograms are shown in the corresponding right panel (b,d). Right panels (b,d): Enrichment histograms showing the fraction of raw reads contributed by each of the 1135 samples to the two genomes whose GC-coverage plots are displayed in the corresponding left panel. x-axis: read abundance of genome bin 470 (panel b) and 766 (panel d); y-axis: sample frequency (among the 1135 samples).

The fact that many of the partitions display low contamination is somehow balanced by the concomitant generation of large and unresolved partitions. The production of these unresolved partitions arises from the fact that the extent of genome divergence is not uniform across the range of taxa occurring in the samples. As discussed above, strain-level (“pangenomic”) variation is another factor contributing to cluster fragmentation, by inducing a differential segregation of the core and variable portions of genomes, and is exacerbated by the inter-sample read aggregation process.

### 2.6 Recovery of very low-abundance genomes

A key motivation for the pre-assembly processing of reads was the theoretical possibility to aggregate reads from low abundance organisms across samples.

To assess whether we could indeed identify such consistently low abundance genomes in real-life datasets, we characterized the abundance of a subset of > 70% complete genomes from the LifeLinesDeep cohort analysis by directly mapping the raw reads of the original samples against them. Given the large size of the cohort, this analysis was not performed on the full set of MAGs but on a limited number of genomes, the aim being to validate the ability of the method to retrieve such low-abundance genomes by exhibiting some of them.

The relative enrichment levels of these genomes was measured as the fraction of raw reads contributed by each sample to them (Methods), and is illustrated in Figure 4 for two genomes, with the left panel showing an example of a consistently low-abundance genome (i.e. with nearly all the samples contributing no more than 10^−5^ to 10^−4^ of their reads to the given genome), while the right panel shows a genome of overall moderate abundance (10^−4^) but reaching higher abundance (10^−3^) in a few dozens of samples (represented by the rightmost peak in the histogram).

Given the large number of microbiomes analyzed, we quite frequently observed situations where a given genome reaches medium to high relative abundance in at least one sample (as illustrated in the right panel of Figure 4). However and importantly, we could also detect instances of genomes that consistently segregated at low abundance levels across the whole cohort (left panel of Figure 4 and right panels of Figure 5).

The recovery of these genomes was made possible by aggregating a few thousands reads per sample across a large number of samples, thus demonstrating the ability of the method to isolate rarer genomes. Overall, the high proportion of homogeneous partitions corresponding to partial genomes (Table 3) is consistent with the recovery of sequences from lower abundance organisms, whose cumulative coverage across the cohort is not sufficient to allow complete genome reconstruction.

### 2.7 Assessing novelty against reference genome compendia

To investigate the extent to which the recovered genomes could correspond to novel organisms, we screened a subset of 164 of them (more than 50% complete with less than 5% contamination, accessible at http://www.genoscope.cns.fr/SCdata/MAGs/) against several reference genome libraries. We first compared the genomes against the Kraken 2 [39] database built from NCBI’s Refseq bacteria, archaea and viral libraries (on October 2018). Only 21 out of the 164 genomes compared had at least one fragment classified against this reference database (Methods). We also compared the genomes against the “Global Human Gastrointestinal Bacteria Genome Collection” (HGG, ref [13]), that represents one of the most comprehensive resources of gastrointestinal bacterial reference sequences currently available. Only less than half (72/164) of the genomes displayed convincing similarity to the HGG genome catalogue (Methods).

## 3 Discussion

Covariance-based binning has the power to identify biologically meaningful associations between metagenomic sequences that could go unnoticed by analyses based on sequence overlap (assembly) or nucleotide signatures. This is illustrated in the present study by a preliminary experiment using a synthetic dataset spiked with low abundance sequences from a target genome that does not reach a sufficient coverage to yield kilobase-sized fragments after assembly in any individual sample (thus precluding the application of contig binning), but which is successfully recovered via read-level binning (Supplementary Table). When applied to the > 10^10^ reads from the LifeLinesDeep cohort’s metagenomes, our bin-first protocol recovers hundreds of metagenome-derived genomes, including from consistently less abundant organisms (Figure 4 and right panels of Figure 5). By increasing the number of distinct abundance profiles that can be generated, larger sample numbers increase both the sensitivity and resolution of covariance-based methods; one may therefore anticipate further gains in the application of such methods in relation to future increases in the scale of sequence data generated (e.g. increased cohort sizes).

We need however to acknowledge several important limitations that impede the overall performance and applicability of our bin-first framework. First, we already mentioned a limitation arising from the natural occurrence of strain-level variation at the origin of differential segregation of core and variable fractions of species pangenomes (Figure 2). The large number of incomplete but otherwise uncontaminated partitions/bins in the LifeLlinesDeep analysis partly reflects the widespread occurrence of this type of variation in natural habitats. However, it should be noted that assembly-based approaches are neither immune to this type of variation, frequently discarding it when building “flattened” consensus contigs. This type of polymorphism is difficult to handle in a *de novo* way, and current methods for strain-level surveys of metagenomes typically rely on reference databases of strain-specific nucleotide polymorphisms (see for example ref [24]). Sample by sample assembly limits the risk of strain mix-up, but at the expense of focusing on those genomes reaching high-coverage (around 10x). Our approach aimed at relaxing the latter constraint, but by doing so through the aggregation of lower abundance reads across samples, it becomes vulnerable to extensive strain-level variation.

To the best of our knowledge, a method that could target -in an unsupervised way-low-coverage genomes in a strain resolved manner is not available today, and working towards this goal is clearly a promising research area. It should be noted however that, to some extent, the degree of similarity that one wishes to distinguish can be tuned through the choice of the k-mer length and hash size. Increasing the k-mer size would increase the separation of closely related sequences, but only to some extent because the locality sensitive hashing scheme will inherently increase the probability of collision for similar sequences. Thus, we face here another benefit *versus* disadvantage trade-off: besides efficient in-memory indexing, LSH allows to conveniently handle sequencing errors (noise), but at the same time can also put a limit on the power to separate very similar sequences (e.g. strains).

The observation that four out of seven genomes retrieved in the preliminary experiment based on 18 samples were not among the set of MAGs identified by analyzing the full dataset is indicative of a lack of stability of the algorithm. This effect of the sample number is most likely mediated by the increasing presence of strain variation when aggregating reads across increasing numbers of samples, leading to more fragmented partitions, and suggests that, above a certain level, increases in sample number can lead to diminishing returns in terms of complete genome recovery. We probably underestimated the extent of strain-level variation in real-world data, and the high-level of genome fragmentation in the LifeLinesDeep partitions can be partly attributed to this problem, with low sequence coverage able to contribute as well.

Another limitation of the method is the generation of coarse-grained partitions harboring a large number of unresolved genomes (corresponding to the tail of the partition size distribution shown in Figure 3). This problem is already manifest in the preliminary experiment comparing assembly-first versus bin-first approaches, and further exemplified in the large cohort analysis that yielded 983 partitions, 888 of which displayed low-levels (<5%) of contamination (Table 3), but also produced several large clusters holding dozens of microbial genomes. The generation of such unresolved partitions appears difficult to avoid as the extent to which genomes differ to each other is variable across phylogenetic groups. As a result, it is unlikely that a single setting (e.g. k-mer length and hash size) could achieve perfect separation of genomes from highly diverse genome mixtures.

These two limitations probably concur to explain that the number of moderate to nearly complete genomes recovered from the population cohort analysis appears much lower than the number of “species genomes” recoverable via assembly-first approaches (remember for example that close to 5,000 species-level genome bins were recovered from the analysis of nearly 10,000 metagenomes in ref [31] (one should however note that an average of 5.3 Gb per sample ^*^after^*^ quality control was generated in the latter study, versus 3.0 Gb ^*^before^*^ quality control in the LLDeep [43] data analyzed here). When analyzing a large number of related samples, we noticed quite commonly that distinct organisms are able to reach a sufficiently high (to be assembled) relative abundance level in at least one sample (a situation exemplified in the right panel of Figure 4). When following a sample by sample assembly-based strategy, a high coverage reached in a single sample (the likelihood of which increases with the number of samples analyzed) might be sufficient to assemble significant portions of a genome, even if it segregates at much lower levels in the remaining part of the cohort. This probably contributes to explain the high genome recovery yields of assembly-based approaches. However, a key feature of the presented method is its ability to recover genomes of organisms consistently segregating at low levels across the entire cohort, as verified in a test experiment and on real-world data (cf left panel of Figure 4). The observation that more than half of the genomes recovered here were not detected in a very large compendium of human gut genomes assembled from thousands of samples [31] is consistent with this view.

Metagenomic sequence binning is still a very active research field, and there are many interesting ongoing efforts, including some attempts to cast binning as an assembly-graph partitioning problem [32]. Recent efforts include ref [6] that exploits structural sparsity of compact De Bruijn assembly graphs to compute succinct indexes in linear time, allowing to perform neighborhood queries on large assembly graphs in an “assembly-free” manner. One should note however that, even though this can leverage developments in efficient k-mer counting and graph compaction (e.g. [10]), assembling large multi-terabytes datasets can remain problematic in the first place. Nevertheless, most of the recent development efforts in the field of metagenomic sequence binning remain directed toward assembly-first approaches, which have already delivered a vast array of performent and user-friendly software [1, 3, 23, 40], some of which have shown capabilities to recover genomes as low as 0.6 % (10^−3^) in relative abundance [40]. However, we have shown that the method presented here is able to recover genomes by sequence enrichments of the order of up to 10^−6^ (10^−7^ for some plasmid sequences), and therefore believe that it could be a useful adjunct to existing more mainstream approaches, especially for targeting more rare organisms. On the other hand, benefits of read-binning for comparative metagenomics have also recently been reported [36].

## 4 Methods

### 4.1 Control datasets

The dataset described in ref [15] (and accessible from http://www.genoscope.cns.fr/SCdata/vc50/) corresponding to a virtual cohort of 50 individuals each harboring a microbiome of 100 distinct bacterial genomes sampled under a power-law abundance distribution (with power parameter *α* = 1.0) from a pool of 750 fully-sequenced genomes at an average depth of 10x (see ref [15] for details), was used in the control experiments. We call these datasets semi-synthetic because they are made of real bacterial genome sequences assembled into artificial mixtures. The read to genome assignments (ground truth) being known in advance for all the reads, the precision and recall metrics were computed from the read clustering output as in equations (10) and (11) from ref [27] (see section “Comparison of read binning algorithms”), with precision corresponding to what the authors refer to as purity and recall corresponding to completeness.

### 4.2 Real dataset: LifeLines-DEEP metagenomes

The LifeLines-DEEP cohort features 1135 individuals (474 males and 661 females) from the general Dutch population, whose gut microbiomes were shotgun sequenced using the Illumina short read technology, generating an average of 32 million reads per sample, see [43] and EBI dataset accession number EGAD00001001991.

### 4.3 Locality Sensitive Hashing (LSH)

We used the SimHash [8] scheme described in ref [11] to obtain a proxy for k-mer abundance. Briefly, raw reads are parsed into k-mers of fixed size (k=31 was used in our experiments), the bases of which are individually mapped to a complex simplex via a mapping of the form: A=1, C=i, G=-i, T=-1, that can also incorporate base-call confidence scores [11]. k-mers are thus represented in k-dimensional space in which *n* hyperplanes (we used *n* = 30 in our experiments) are randomly drawn, creating 2^*n*^ subspaces, or buckets, indexing the columns of the sample by k-mer abundance matrix whose rows were scaled to unit *𝓁*_2_ norm. The LSH scheme is sequence sensitive, increasing the probability of collision for more similar k-mers [8], and allows the representation of k-mer abundance matrices of arbitrary dimensions in fixed memory.

Regarding the choice of a k-mer length, the key requirement is that k-mers should be sufficiently long so that most of them will be specific to each genome, thereby capturing genuine abundance patterns of individual genomes. In our experiments, the k-mer length (31) was chosen to be close to the value used by the authors of ref [11] to analyze their largest (terabase-sized) dataset. Some limited experiments with varying k-mer length values were carried out on smaller subsets of the data to check that small variations in k-mer size did not result in disproportionate differences in clustering outputs. In choosing the k-mer length, we were also guided by the observations in ref [21] that k-mer similarity between genomes at different k approximates various degrees of taxonomic similarity, with k=31 appearing to correspond to species-level similarity. We also noticed that k=31 is the default setting in the popular sequence classification engine kraken [39].

### 4.4 Sparse coding

Our aim is to learn sparse and non negative factors from the sample by (hashed) k-mer abundance matrix **X**. The sparsity assumption has biological roots in the fact that every individual only harbors a small subset of all the genomes that constitute the global microbiome, while each genome only contains a very small subset of the k-mers encountered across all the samples. Sparse coding aims at modeling data vectors as sparse linear combinations of elements of a basis set (aka dictionary) that can be learned from the data by solving an optimization problem [25]. We used the spams library (http://spams-devel.gforge.inria.fr/), that implements the learning algorithm of ref [25]: given a training set **x**^1^, …, **x**^*n*^ it tries to solve

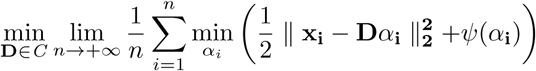

where *ψ* is a sparsity-inducing regularizer (e.g. the *𝓁*_1_ norm) and *C* is a constraint set for the dictionary (positivity constraints can be added to *α* as well). The following optimization scheme was used (FL stands for fused lasso):

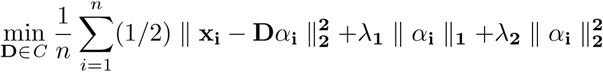

with *C* a convex set verifying

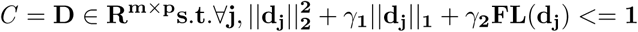

Once the dictionary has been learned, the spams library offers an efficient implementation of the LARS algorithm [12] for solving the Lasso or Elastic-Net problem: given the data matrix **X** in **R**^*m×n*^ and a dictionary **D** in **R**^*m×p*^, this algorithm returns a matrix of coefficients **A** = [*α*^**1**^, …, *α*^**n**^] in **R**^*p×n*^ such that for every column **x** of **X**, the corresponding column *α* of **A** is the solution of

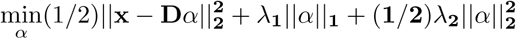

The spams implementation of this algorithm allows to add positivity constraints on the solutions *α*, which have a natural interpretation as weighing the contribution of the different hashed k-mers to the latent genomes. In practice, we defined clusters by assigning hashed k-mers from bucket *i* to component *c* if *c* = argmax_*j*_ *A*_*i,j*_.

### 4.5 Read classification and Assembly

Starting with the raw reads and their decomposition into k-mers, the bulk of the binning algorithm thus operates in k-mer space. After computing co-varying k-mers sets (“eigengenomes”), a post-processing step is thus necessary to assign reads to their cognate k-mer clusters in order to achieve a read-level clustering. We sticked to the LSA procedure [11] for this step, with the original reads being assigned to k-mer clusters based on a log-likelihood score aggregating i) cluster sizes (measured in terms of k-mer numbers), ii) the overlap between k-mers in reads and those in clusters, iii) an inverse document frequency (IDF)-style weight expressing the rarity of each of the overlapping k-mers. After read assignment, the partitions were assembled with the Spades (v3.13.0) engine [4] using default settings.

### 4.6 First experiment for comparing assembly-first versus bin-first protocols

An experimental setup was designed to illustrate the ability of read binning to cluster rare reads form a target genome across samples, while assembly-first protocols are inoperable because the low coverage of the target genome prevents the generation of any kilobase-sized contig from the assembly of the individual samples. The dataset consisted in 18 samples each containing a subset of 20000 reads sampled from the 18 metagenomic libraries analyzed in ref [34] and randomly spiked with mock reads from a Bacillus thuringiensis plasmid (NG–035027.1) as in the test data used in the original LSA paper [11]. However, as the number of spiked reads (up to 4000) distributed among the samples in LSA’s test dataset was sufficient to yield contigs covering a large fraction of the plasmid genome upon assembly, we derived a new dataset only containing 0 to 100 paired-reads (14 samples contained 100 paired-reads while 4 were entirely devoid of plasmid reads) and used the latter for this experiment. This dataset is available on the website associated to this publication’s material.

After checking that no kilobase-sized contig could be assembled in any of the samples -thus precluding the application of contig binning-the dataset was processed by our pre-assembly pipeline using the following settings: a k-mer length of 30 and a hash size of 22 were used to build the k-mer abundance matrix, the latter was decomposed by a svd and the columns of the eigen-kmer matrix were clustered using a cosine similarity threshold of 0.25, followed by read assignment and assembly (using Spades) of the partitions. More than 99% (2782 out of 2800) of the plasmid derived reads ended up in a single partition (Supplementary Table), leading to the recovery of the complete target genome sequence upon assembly.

### 4.7 Second experiment for comparing assembly-first versus bin-first strategies

The raw sequence data from 18 (randomly chosen) individuals of the LLDeep cohort were either assembled individually (i.e. on a sample by sample basis) with metaSPAdes (v3.13.0) followed by contig binning across samples with the MetaBat2 adaptive algorithm [19], or the raw reads were clustered using our read-level binning pipeline, followed by metaSPAdes assembly of the resulting partitions/bins.

The raw reads were first mapped to the assembled contigs using bwa-mem [22] using default parameters. MetaBat2 was then invoked in the following way: first, the jgi summarize bam contig depths script was called to compute contig abundance statistics from the read mapping bam files, with the default options (minimum percent identity for a mapped read: 0.97, minimum contig length: 1000, minimum contig depth: 1). The metabat2 program was then called using the default parameters (minCV 1.0, minCVSum 1.0, maxP 95%, minS 60, and maxEdges 200) and the previously generated coverage statistics file, leading to the generation of 225 bins covering 694000907 bases.

For the comparison, our sparse coding pipeline was then executed under the same settings as in the full cohort analysis (hash size and k-mer size equal to 30 and 31 respectively, and default parameters for the dictionary learning and sparse decomposition of the abundance matrix), with the exception of the number of components that was matched to the number of bins (225) generated by MetaBat2. To generate Figure 1, the complete genomes retrieved using both approaches were aligned (using nucmer [26] with default parameters) to individual assemblies from all the samples, and the number of distinct contig hits (≥99% identity and ≥ 2500 bp) was recorded.

### 4.8 Comparison of read binning algorithms

The virtual cohort dataset described above and in ref [15] was used to compare the clustering accuracies of the original LSA [11] and sparse coding methods, as well as the performance of directly clustering the columns of the abundance matrix using a k-means algorithm as a baseline. The reads to genome memberships being comprehensively known in these controlled genome mixtures, clustering accuracy metrics (precision, recall and F-measure) could be quantified as in ref [27] (Table 1). Briefly, each bin is first mapped to its most abundant (in terms of number of reads) genome (note that if each bin is mapped to a single genome, a given genome can be mapped to multiple bins). Precision is defined as the ratio of reads originating from the mapped genome to all the bin’s reads. Recall on the other hand reflects how complete a bin is with respect to the sequence of its cognate (mapped) genome. Average precision is the fraction of correctly assigned reads for all assignments to a given cluster averaged over all clusters, while average completeness is averaged over all genomes (including those possibly not assigned to any cluster). We follow ref [27] in order to give larger bins higher weight in performance determinations. Specifically, if *X* is the set of clusters and *Y* the set of underlying genomes, precision and recall are defined respectively as:

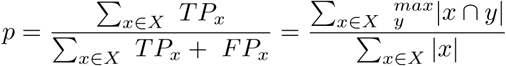

and

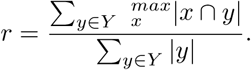

The same k-mer abundance matrices (built using a k-mer size of 31 and a number of hash bits (hyperplanes) equal to 30) were used as input for all the methods.

### 4.9 Initial estimate of genome richness and number of components

For the test experiments based on synthetic microbiomes of controlled complexity (e.g. the virtual cohort of 50 individuals, where each microbiome consisted in 100 genomes drawn from a pool of 750 genomes under a given abundance distribution), the number of clusters was set to match the (known) number of distinct genomes segregating in the complete set of samples. For the analysis of real-world data (i.e. the LifeLinesDeep cohort), where the total number of genotypes is unknown, a meaningful number of components for the sparse decomposition was estimated on the basis of the number of distinct rpS3 ribosomal protein sequences in the analyzed metagenomes, clustered at 98% identity, which roughly corresponds to species level delineations according to ref [35].

### 4.10 Evaluation of read enrichment levels

To assess whether we could identify genomes segregating at consistently low abundance levels in real-life datasets, we characterized the abundance of a dozen of MAGs reconstructed from the LifeLinesDeep cohort analysis by directly mapping the raw reads from the original samples against them. Given the large size of the cohort (and the significant amount of computer resources associated with this analysis), and given that our objective was to establish whether consistently rare genomes can be identified by the method or not, this analysis was performed on a limited number of genomes.

Relative enrichment levels were estimated by mapping the original reads (after removal of duplicated reads) to the genome-resolved partitions using bwamem [22] with default parameters. Uniquely and consistently (i.e. paired) mapped reads were scored to compute enrichment ratios as the number of mapped reads divided by the number of raw reads analyzed, as displayed for example on the x-axes of Figure 4 and the right panels of Figure 5.

### 4.11 Comparison of genome-resolved partitions to reference genomes

To assess the novelty of the genomes assembled from individual partitions produced by our pipeline through the analysis of the LifeLines-Deep cohort, we screened them against two reference libraries. First, the genomes were compared to the Kraken2 (v1) database (https://ccb.jhu.edu/software/kraken2/) built from NCBI’s refseq bacteria, archaea, and viral libraries (in August 2018), using the Kraken2 classifier [39] and a confidence score threshold of 0.2. Second, the same genomes were compared against the Human Gastrointestinal Bacteria Genome Collection [13] (HGG, encompassing more than 100 GB of sequence data) using the nucmer aligner [26] with default parameters. A genome was marked as already known if it shared at least ten distinct 99% identity alignments of length ≥5 kb to any HGG entry.

### 4.12 Binning implementation

Code for the pipeline used to perform the analysis of the LifeLinesDeep cohort can be cloned from https://gitlab.com/kyrgyzov/lsa_slurm, while a more lightweight implementation of key algorithms (including sparse NMF) can be downloaded from https://github.com/vincentprost/LSA_NMF; they draw on the code base of the LSA tool (ref [11] and https://github.com/brian-cleary/LatentStrainAnalysis), and on the SPAMS (SPArse Modeling Software) library that can be downloaded from http://spams-devel.gforge.inria.fr/. The analysis of the metagenomes from the LifeLines-DEEP cohort was carried out on a Bullion S6130 octo module server equipped with 2 Intel Xeon Haswell E7-4890 v3 CPU (18 cores) per module, 8 TB of RAM and 35 TB storage. Most of the tasks being embarassingly parallelizable, they were run through a Slurm workload manager. The analysis took about three weeks wall time, with the sparse decomposition of the k-mer abundance matrix taking less than one day. The bulk of the execution time was spent in pre and post-processing tasks: pre-processing of the 10 terabytes of raw reads to improve load balancing (∼5 days), k-mer hashing and counting for constructing the k-mer abundance matrix (∼4.5 days), assignments of reads to eigengenomes following the sparse decomposition step (∼ 6 days), and assembly of individual read partitions using the spades assembly engine [4] (∼2.5 days).

A desirable feature when designing computational pipelines is to have resource requirements, especially memory, scale in a way independent of the sheer data volume. This is the case for the analytical method presented here, as it can be executed “in memory” with the dimensionality of the empirical abundance matrix tailored via the LSH scheme to capture the desired amount of sequence diversity while remaining consistent with the available resource budget. The use of efficient online matrix factorization techniques [25] leads to limited memory footprints. Even though we leveraged here a powerful computer infrastructure to carry out the analysis of the large cohort dataset (10 terabytes of data), our pipeline is routinely executed on commodity hardware for smaller projects.

## 5 Availability of source code

Code for the pipeline used in the analysis of the LifeLinesDeep cohort can be cloned from https://gitlab.com/kyrgyzov/lsa_slurm, while a more lightweight implementation of key algorithms (including sparse NMF) can be downloaded from https://github.com/vincentprost/LSA_NMF.

## 6 Availability of supporting data and materials

Assembled sequences of the genome-resolved bins (more than 50% complete and with less than 5% contamination) recovered from the analysis of the LifeLines-DEEP cohort are available at http://www.genoscope.cns.fr/SCdata/MAGs/. The dataset corresponding to the virtual cohort used in the test experiments can be downloaded from http://www.genoscope.cns.fr/SCdata/vc50/, and the reduced spiked dataset used in the preliminary experiment from http://www.genoscope.cns.fr/SCdata/benchs/spike/.

